# Coffee induces chromatin structural changes associated with histone H4K16 hypoacetylation

**DOI:** 10.64898/2026.10.13.758977

**Authors:** Yasuko Tanaka, Yangsu Cho, Yuusuke Suwabe, Sora Misaki, Ryo Uchidaya, Kazumasa Aoyama

**Affiliations:** Division of Hygienic Chemistry, Faculty of Pharmacy, Keio University, Tokyo, Japan

**Keywords:** Coffee, H4K16ac, Histone acetylation, Chromatin structure, Gene expression

## Abstract

Histone acetylation plays a critical role in the regulation of chromatin structure and gene expression. Among histone acetylation marks, acetylation of histone H4 at lysine 16 (H4K16ac) is particularly important for the maintenance of chromatin organization. We previously identified a reduction in H4K16ac following coffee treatment through mass spectrometry-based histone modification profiling. In the present study, we investigated the effects of coffee on H4K16ac levels, chromatin structure, and gene expression. Western blotting and immunofluorescence analyses confirmed that coffee treatment reduced H4K16ac levels in a cell type-dependent manner. Notably, H4K16ac was reduced in HCT116 cells, in which H3K27ac was not decreased by coffee treatment. Quantitative analysis of DAPI-stained nuclei revealed a significant increase in the coefficient of variation of nuclear fluorescence intensity following coffee treatment, indicating alterations in chromatin structure. Furthermore, integrative analysis of publicly available H4K16ac chromatin immunoprecipitation sequencing (ChIP-seq) data and previously generated RNA sequencing (RNA-seq) data revealed that H4K16ac-enriched genes exhibited transcriptional changes following coffee treatment in K562 cells, including the downregulation of MYC-associated genes. Together, these findings demonstrate that coffee treatment induces alterations in chromatin structure accompanied by H4K16 hypoacetylation and suggest a potential association between these epigenetic changes and transcriptional regulation.

## Introduction

Chromatin structure plays a fundamental role in the regulation of gene expression, DNA replication, and genome stability. The organization of chromatin is dynamically regulated by post-translational modifications of histones, including acetylation, methylation, phosphorylation, and ubiquitination (Bannister & Kouzarides, 2011; Kouzarides, 2007). Among these modifications, histone acetylation is generally associated with increased chromatin accessibility and transcriptional activity. Acetylation of lysine residues neutralizes their positive charge and can influence histone–DNA interactions, nucleosome organization, and the recruitment of chromatin-associated proteins (Bannister & Kouzarides, 2011; Roth *et al*, 2001).

Acetylation of histone H4 at lysine 16 (H4K16ac) is particularly important for the regulation of higher-order chromatin structure. Unlike many other histone acetylation marks, H4K16ac directly affects interactions between neighboring nucleosomes and modulates chromatin compaction. Shogren-Knaak et al. demonstrated that H4K16 acetylation inhibits the formation of compact chromatin fibers and influences interactions between chromatin and non-histone proteins (Shogren-Knaak *et al*, 2006). Subsequent studies further established the importance of H4K16ac in regulating chromatin fiber organization(Robinson *et al*, 2008). These findings indicate that changes in H4K16ac levels may have substantial effects on chromatin architecture.

Alterations in H4K16ac have also been implicated in cellular processes associated with transcriptional regulation, genome maintenance, and disease. Loss of H4K16ac has been observed in several human cancers, suggesting that dysregulation of this modification may contribute to pathological changes in chromatin organization (Fraga *et al*, 2005). Furthermore, we previously demonstrated that nuclear c-Abl-mediated signaling induces chromatin structural changes accompanied by H4K16 hypoacetylation (Aoyama *et al*, 2011). These observations suggest that H4K16ac may serve as an important link between cellular signaling pathways and chromatin organization.

Coffee contains numerous bioactive compounds and has been reported to influence a variety of cellular processes. In our previous study, we demonstrated that coffee treatment induces widespread changes in histone acetylation and transcriptional regulation, including suppression of MYC-associated transcriptional programs and inhibition of cell proliferation (Cho *et al*, 2026). Mass spectrometry-based histone modification profiling further revealed a reduction in H4K16ac following coffee treatment. However, whether coffee-induced H4K16 hypoacetylation is associated with alterations in chromatin structure remains unknown. Furthermore, the potential relationship between H4K16ac enrichment and coffee-induced transcriptional changes has not been investigated.

In the present study, we investigated the effects of coffee treatment on H4K16ac levels, nuclear chromatin structure, and gene expression. We first examined coffee-induced changes in H4K16ac across multiple cell types using Western blotting and immunofluorescence analyses. We then evaluated chromatin structural changes in HCT116 cells by quantifying the spatial heterogeneity of nuclear DNA staining intensity. Finally, we integrated publicly available H4K16ac chromatin immunoprecipitation sequencing (ChIP-seq) data with our previously generated RNA sequencing (RNA-seq) data to investigate the relationship between H4K16ac enrichment and coffee-induced transcriptional repression, with particular emphasis on MYC-associated genes. Through these analyses, we sought to clarify the potential relationship between coffee-induced H4K16 hypoacetylation, chromatin organization, and transcriptional regulation.

## Materials and Methods

### Cell culture

K562, HCT116, A431, MCF7, A549, and TIG-1 cells were cultured and maintained as previously described (Cho *et al*., 2026; Higashi *et al*, 2026). Coffee was prepared according to the procedure described in our previous study (Cho *et al*., 2026). Cells were treated with 5% (v/v) coffee for 24 h.

### Antibodies

The following antibodies were used in this study: anti-H4K16ac (#13534, Cell Signaling Technology), anti-β-actin (#sc-47778, Santa Cruz Biotechnology), anti-histone H2A (#ab18255, Abcam), HRP-conjugated anti-mouse IgG (light chain-specific, #115-035-174, Jackson ImmunoResearch), HRP-conjugated anti-rabbit IgG (light chain-specific, #211-032-171, Jackson ImmunoResearch), and Alexa Fluor 488-conjugated anti-rabbit IgG (#ab150077, Abcam).

### Western blot analysis

Western blot analysis was performed as previously described(Aoyama *et al*, 2018; Aoyama *et al*, 2021; Aoyama *et al*, 2015). Briefly, cells were harvested following coffee treatment, and protein extracts were subjected to SDS-PAGE, followed by transfer onto membranes. Membranes were incubated with the indicated primary antibodies, followed by appropriate HRP-conjugated secondary antibodies. Protein bands were detected and quantified by densitometric analysis as previously described (Mashimo *et al*, 2019; Yuki *et al*, 2015; Yuki *et al*, 2019). H4K16ac signals were normalized to histone H2A in Fig. 1B and to β-actin in Fig. 1C and F. Relative H4K16ac levels were calculated by setting the corresponding untreated control to 1.

**Figure 1.**
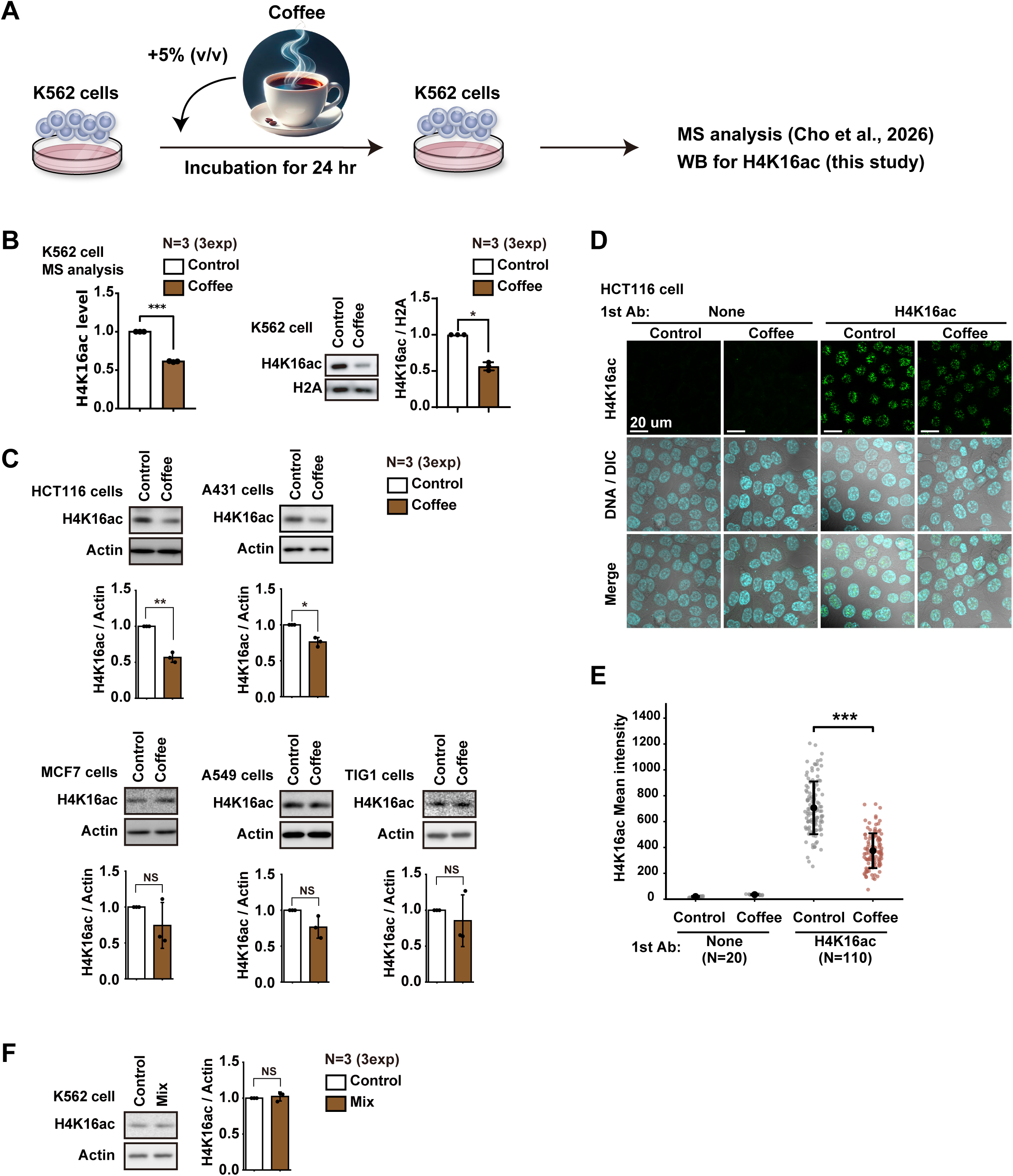
Coffee induces H4K16 hypoacetylation in a cell type-dependent manner. (A) Schematic overview of the experimental strategy used to investigate coffee-induced changes in H4K16ac. Histone modification profiling by mass spectrometry was performed in K562 cells treated with 5% (v/v) coffee for 24 h, as previously reported (Cho et al., 2026). H4K16ac levels were subsequently examined by Western blotting and immunofluorescence analyses. (B) Analysis of H4K16ac levels in K562 cells following coffee treatment. Mass spectrometry data from Cho et al. (2026) and representative Western blot images are shown. H4K16ac signals were normalized to histone H2A. (C) Western blot analysis of H4K16ac levels in HCT116, A431, MCF7, A549, and TIG-1 cells following coffee treatment. H4K16ac signals were normalized to β-actin, and relative levels were calculated by setting the corresponding untreated control to 1. (D) Representative confocal immunofluorescence images of H4K16ac (green) and DAPI (blue) in control and coffee-treated HCT116 cells. Negative controls were prepared by omitting the primary antibody. (E) Quantification of nuclear H4K16ac fluorescence intensity in HCT116 cells. A total of 110 cells per condition were analyzed from two independent biological replicates (55 cells per condition per replicate). Coffee treatment reduced H4K16ac fluorescence intensity by approximately 47% in both replicates. Statistical comparisons were performed separately for each replicate using two-tailed Welch’s t-tests on individual cell measurements (rep1, p = 4.14 × 10 ¹ ; rep2, p = 3.39 × 10 ²). (F) Western blot analysis of H4K16ac levels in K562 cells treated with a mixture of selected coffee constituents, as described previously (Cho et al., 2026). H4K16ac signals were normalized to β-actin. Western blot experiments were performed using three independent biological replicates. Statistical significance was defined as p < 0.05.

### Immunofluorescence staining and microscopy

Immunofluorescence staining and confocal microscopy were performed as previously described (Cho *et al*., 2026; Higashi *et al*, 2025; Taguchi *et al*, 2026). HCT116 cells were seeded onto poly-L-lysine-coated glass coverslips and treated with 5% (v/v) coffee for 24 h. Cells were fixed with 4% paraformaldehyde for 20 min at room temperature, washed three times with phosphate-buffered saline (PBS), and permeabilized and blocked with PBS containing 0.5% Triton X-100, 1% bovine serum albumin, and 0.02% sodium azide for 20 min. Cells were then incubated with an anti-H4K16ac antibody (1:200), followed by an Alexa Fluor 488-conjugated anti-rabbit IgG secondary antibody (1:1,000). After washing with PBS, coverslips were mounted using VECTASHIELD mounting medium containing DAPI. Negative controls were prepared by omitting the primary antibody.

Fluorescence images were acquired using an FV4000 laser scanning confocal microscope (Evident, Japan). Quantification of nuclear H4K16ac fluorescence intensity was performed using ImageJ, based on the method described by our previous papers (Aoyama *et al*., 2011; Aoyama *et al*., 2015; Aoyama *et al*, 2013). Nuclear regions of interest were defined using DAPI staining, and the mean H4K16ac fluorescence intensity within each nucleus was measured.

### Quantitative analysis of chromatin structure

Chromatin structural changes were evaluated by quantitative analysis of nuclear DNA staining intensity, based on previously described methods (Aoyama *et al*., 2011; Kubota *et al*, 2013; Kubota *et al*, 2015; Takahashi *et al*, 2009), with modifications. Nuclear regions of interest (ROIs) were defined based on DAPI staining, and the fluorescence intensity of individual pixels within each nucleus was measured using ImageJ. The mean fluorescence intensity and standard deviation (SD) of pixel intensities were calculated for each nucleus. To account for differences in mean fluorescence intensity among individual nuclei, the coefficient of variation (CV) was calculated as SD divided by the mean fluorescence intensity. Whereas previous studies evaluated chromatin structural changes using the SD of nuclear DNA staining intensity, the present study employed CV to normalize fluorescence intensity variability to its mean.

### ChIP-seq data analysis

Publicly available H4K16ac ChIP-seq data from K562 cells (GSE158736; GSM4809274) were obtained from the Gene Expression Omnibus (GEO) database. The processed H4K16ac signal track (GSM4809274_K562_H4K16ac.bw) was used for subsequent analyses. The distribution of H4K16ac signals across genes was analyzed using deepTools(Ramírez *et al*, 2016). Signal matrices were generated using computeMatrix scale-regions, covering gene bodies and 5-kb regions upstream and downstream of transcription start and end sites, respectively. Genes were classified into four clusters according to their H4K16ac enrichment patterns using k-means clustering (k = 4) implemented in plotHeatmap. Heatmaps and average signal profiles were generated to visualize H4K16ac enrichment patterns (Fig. 3A). To examine the association between H4K16ac enrichment and gene expression, genes in each cluster were compared with RNA-seq data from coffee-treated K562 cells, as described below (Fig. 3B, C). For visualization of individual genomic loci, H4K16ac signal tracks were examined at the MCM7, HNRNPA3, CDCA7, and NME2 loci (Fig. 4B). Mean H4K16ac signals across the gene bodies of these genes were calculated and compared with the genome-wide mean signal (Fig. 4C).

### RNA-seq data analysis

RNA-seq data from control and coffee-treated K562 cells were obtained from our previous study (Cho *et al*., 2026). The RNA-seq data from two independent biological replicates (rep1 and rep2) were reanalyzed to investigate the relationship between H4K16ac enrichment and coffee-induced transcriptional changes. Gene expression levels were quantified as reads per million (RPM). For each replicate, expression fold changes were calculated by comparing coffee-treated cells with their corresponding untreated controls and expressed as log2 fold changes (log2FC). Genes were assigned to four clusters based on the H4K16ac ChIP-seq analysis described above, and the distributions of log2FC values were compared among clusters (Fig. 3B). To identify genes consistently downregulated by coffee treatment, genes exhibiting a greater than 1.5-fold reduction in expression were selected separately for rep1 and rep2. The overlap between downregulated genes from both replicates was then determined among genes belonging to H4K16ac Cluster 1 (Fig. 3C). Expression levels of selected MYC-associated genes, including MCM7, HNRNPA3, CDCA7/JPO1, and NME2, were extracted from the RNA-seq datasets and presented as RPM values for control and coffee-treated samples (Fig. 4D).

### Gene Ontology enrichment analysis

Gene Ontology (GO) enrichment analysis was performed using g:Profiler(Raudvere *et al*, 2019) to characterize the biological functions of genes associated with H4K16ac enrichment and transcriptional repression following coffee treatment. The analysis was conducted using the 33 genes belonging to H4K16ac Cluster 1 that exhibited greater than 1.5-fold downregulation in both RNA-seq replicates (Fig. 3C). Enrichment analysis was performed for GO biological process (GO:BP) terms. The enriched terms were visualized as bar graphs using the negative logarithm of the nominal p-value (−log10 p-value) (Fig. 3D).

### Statistical analysis

Statistical analyses were performed to evaluate differences between control and coffee-treated cells. Data are presented as the mean ± standard deviation (SD), unless otherwise indicated. Comparisons between two groups were performed using two-tailed Student’s t-tests, except where otherwise specified. Statistical significance was defined as p < 0.05. Western blot analyses were performed using three independent biological replicates (Fig. 1B, C, F). Immunofluorescence and chromatin structure analyses were performed using two independent biological replicates (Figs. 1E and 2). Cells were independently cultured and treated with coffee, followed by simultaneous fixation and staining. Equal numbers of cells were quantified from each biological replicate. For immunofluorescence and chromatin structure analyses, statistical comparisons between control and coffee-treated cells were performed separately for each biological replicate using two-tailed Welch’s t-tests on individual cell measurements. These analyses assessed differences at the cellular level within each replicate, rather than statistical significance across biological replicates. For RNA-seq analyses, differences in the distributions of log2 fold changes among H4K16ac-defined gene clusters were statistically evaluated (Fig. 3B). Nominal p-values were used to indicate statistical significance.

**Figure 2.**
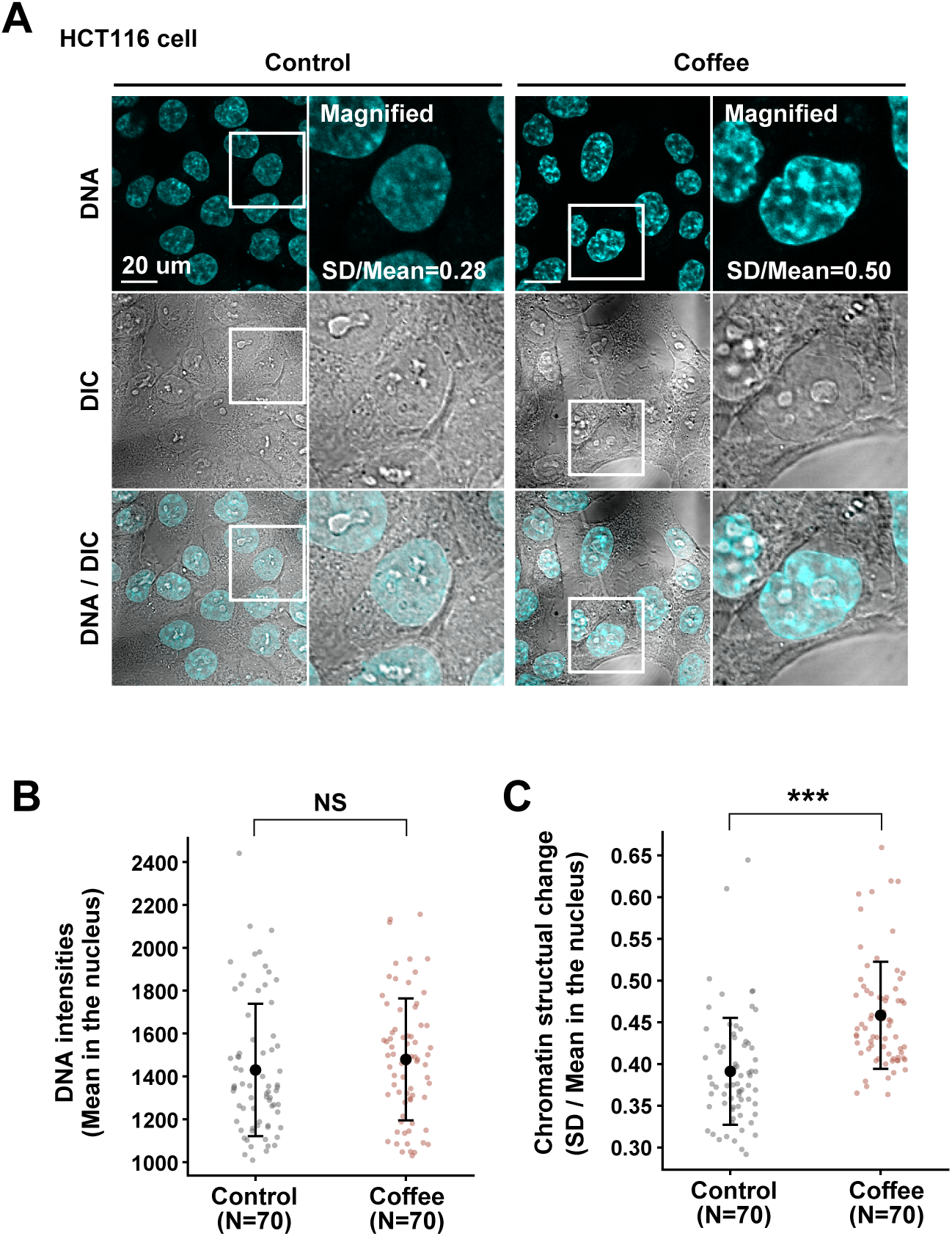
Coffee treatment increases the spatial heterogeneity of nuclear DNA staining intensity in HCT116 cells. (A) Representative confocal images of DAPI-stained nuclei in control and coffee-treated HCT116 cells. Nuclear DNA fluorescence intensity was quantified at the individual pixel level to evaluate changes in the spatial distribution of DNA staining. (B) Quantification of mean nuclear DAPI fluorescence intensity in control and coffee-treated HCT116 cells. No statistically significant difference was observed in either independent biological replicate (rep1, p = 0.256; rep2, p = 0.704). (C) Quantification of the coefficient of variation (CV) of nuclear DAPI fluorescence intensity. CV was calculated for each nucleus by dividing the standard deviation (SD) of pixel fluorescence intensity by its mean. Coffee treatment significantly increased CV in both biological replicates (rep1, +25.5%, p = 6.07 × 10 ; rep2, +9.6%, p = 0.0103). A total of 110 cells per condition were analyzed from two independent biological replicates (55 cells per condition per replicate). Statistical comparisons were performed separately for each replicate using two-tailed Welch’s t-tests on individual cell measurements. These tests evaluate cell-level differences within each replicate.

## Results

### Coffee globally reduces histone H4 lysine 16 acetylation in human cells

In our previous study, mass spectrometry-based histone modification profiling revealed that coffee treatment reduced histone H4 lysine 16 acetylation (H4K16ac) levels (Cho *et al*., 2026). Given the critical role of H4K16ac in chromatin organization, we sought to further characterize the effects of coffee on this histone modification. K562 cells were treated with 5% coffee for 24 h (Fig. 1A). Consistent with our previous mass spectrometry findings, Western blot analysis confirmed a significant reduction in H4K16ac levels following coffee treatment (Fig. 1B). We next examined whether coffee-induced H4K16 hypoacetylation was observed in other cell types. Western blot analysis revealed significant reductions in H4K16ac levels in HCT116 and A431 cells, whereas no significant changes were detected in MCF7, A549, or TIG-1 cells (Fig. 1C). These results indicate that the effects of coffee on H4K16ac vary among cell types. Immunofluorescence analysis further confirmed a reduction in nuclear H4K16ac fluorescence intensity following coffee treatment in HCT116 cells (Fig. 1D, E). Quantitative analysis revealed that H4K16ac fluorescence intensity decreased by approximately 47% in both independent biological replicates. This reduction was statistically significant in each replicate (rep1, p = 4.14 × 10 ¹ ; rep2, p = 3.39 × 10 ² ; Welch’s t-test based on individual cell measurements), demonstrating reproducible H4K16 hypoacetylation following coffee treatment. These findings independently support the results obtained by Western blotting.

In our previous study, we attempted to identify the coffee constituents responsible for its effects on histone acetylation (Cho *et al*., 2026). To determine whether these constituents could also account for the reduction in H4K16ac, we examined the effects of the same mixture in K562 cells. However, the mixture did not significantly reduce H4K16ac levels (Fig. 1F), indicating that the constituents responsible for coffee-induced H4K16 hypoacetylation remain unidentified. Together, these results demonstrate that coffee treatment induces H4K16 hypoacetylation in a cell type-dependent manner, although the specific coffee constituents responsible for this effect remain to be determined.

### Coffee induces alterations in chromatin structure in HCT116 cells

Histone H4K16ac plays an important role in the regulation of higher-order chromatin structure. Given that coffee treatment significantly reduced H4K16ac levels in HCT116 cells (Fig. 1C–E), we investigated whether coffee treatment also affected nuclear chromatin organization. Chromatin structural changes were evaluated by quantitative analysis of the spatial distribution of nuclear DNA fluorescence intensity using DAPI staining, based on previously established methods (Aoyama *et al*., 2011; Takahashi *et al*., 2009). HCT116 cells were treated with coffee, and the fluorescence intensity of individual pixels within each nucleus was quantified (Fig. 2A). We first examined the mean nuclear DAPI fluorescence intensity to assess potential differences in overall DNA staining intensity. No significant difference was observed between control and coffee-treated cells in either of the two independent biological replicates (Fig. 2B). We next assessed the spatial heterogeneity of nuclear DNA staining by calculating the coefficient of variation (CV), defined as the standard deviation (SD) of pixel fluorescence intensity divided by its mean. This normalization allowed us to evaluate fluorescence intensity variability while accounting for differences in mean intensity among individual nuclei. Notably, coffee treatment increased the CV of nuclear DAPI fluorescence intensity in both independent biological replicates (Fig. 2C). The CV increased by 25.5% in rep1 and 9.6% in rep2, with statistically significant differences observed in cell-level comparisons within each replicate (rep1, p = 6.07 × 10 ; rep2, p = 0.0103; Welch’s t-test). Thus, although the overall nuclear DAPI fluorescence intensity remained unchanged, coffee treatment reproducibly increased the spatial heterogeneity of DNA staining. These findings indicate that coffee treatment alters the intranuclear distribution of DNA fluorescence intensity, consistent with changes in chromatin organization. Together with the reduction in H4K16ac levels observed in HCT116 cells (Fig. 1C–E), these results suggest a potential association between coffee-induced H4K16 hypoacetylation and alterations in nuclear chromatin structure.

### H4K16ac-enriched genes exhibit transcriptional changes following coffee treatment

To investigate the potential relationship between H4K16ac and coffee-induced transcriptional changes, we integrated publicly available H4K16ac chromatin immunoprecipitation sequencing (ChIP-seq) data with our previously generated RNA sequencing (RNA-seq) data from coffee-treated K562 cells (Cho *et al*., 2026). We first analyzed the genomic distribution of H4K16ac using the publicly available ChIP-seq dataset and classified genes into four clusters based on their H4K16ac enrichment patterns (Fig. 3A). We then examined coffee-induced transcriptional changes in each cluster using RNA-seq data from two independent replicates (rep1 and rep2). Comparison of gene expression changes revealed that genes in Cluster 1 exhibited significantly lower expression fold changes following coffee treatment than those in Clusters 3 and 4 in both RNA-seq replicates, whereas no significant difference was observed between Clusters 1 and 2 (Fig. 3B). We therefore focused on Cluster 1 and identified genes whose expression was reduced by more than 1.5-fold following coffee treatment in each RNA-seq replicate. Among Cluster 1 genes, 82 and 147 genes were downregulated in rep1 and rep2, respectively. Of these, 33 genes were commonly downregulated in both replicates (Fig. 3C). We next performed Gene Ontology (GO) enrichment analysis of these 33 genes to investigate their biological functions. The analysis revealed enrichment of processes related to nucleic acid metabolism, RNA metabolism, RNA processing, and DNA replication initiation (Fig. 3D). These findings suggest that genes associated with H4K16ac enrichment and consistently downregulated by coffee treatment are involved in fundamental cellular processes, particularly nucleic acid metabolism and RNA-related functions. Together, these results demonstrate that a subset of H4K16ac-enriched genes exhibits reproducible transcriptional downregulation following coffee treatment. Although the publicly available ChIP-seq dataset does not establish coffee-induced changes in H4K16ac occupancy at individual loci, these findings suggest a potential association between H4K16ac enrichment and the transcriptional response to coffee treatment.

**Figure 3.**
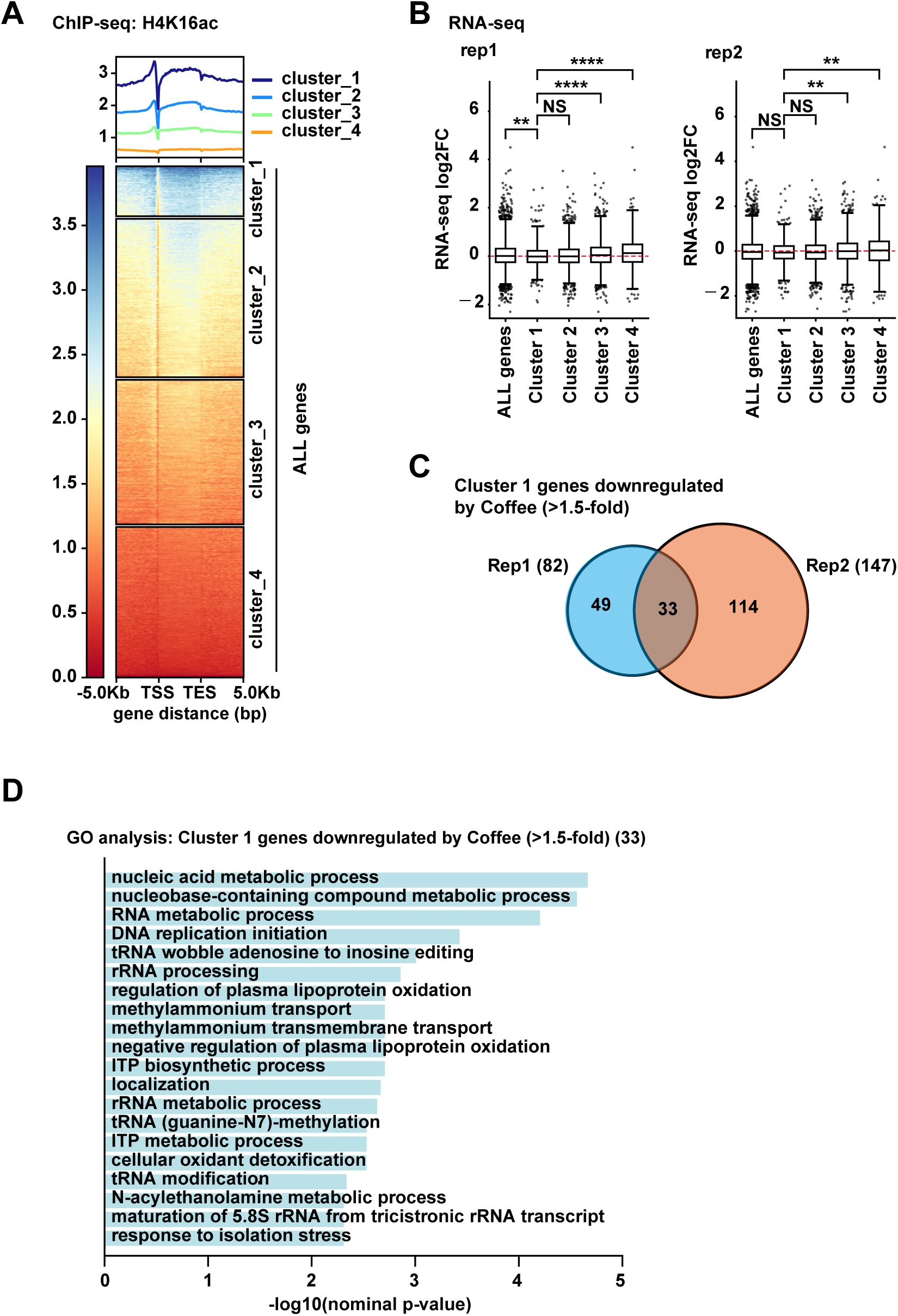
H4K16ac-enriched genes exhibit transcriptional changes following coffee treatment. (A) Heatmap and average signal profiles showing the distribution of H4K16ac across gene bodies and their flanking regions in K562 cells. Publicly available H4K16ac ChIP-seq data (GSE158736; GSM4809274) were analyzed using deepTools. Genes were classified into four clusters by k-means clustering (k = 4): Cluster 1 (n = 1,990), Cluster 2 (n = 6,320), Cluster 3 (n = 5,761), and Cluster 4 (n = 6,004). (B) Comparison of coffee-induced transcriptional changes among the four H4K16ac-defined gene clusters. Log2 fold changes in gene expression were calculated using previously generated RNA-seq data from control and coffee-treated K562 cells (Cho et al., 2026). Results from two independent biological replicates (rep1 and rep2) are shown separately. Cluster 1 exhibited significantly lower expression fold changes than Clusters 3 and 4 in both replicates. (C) Identification of H4K16ac Cluster 1 genes consistently downregulated following coffee treatment. Among Cluster 1 genes, 82 and 147 genes exhibited greater than 1.5-fold downregulation in rep1 and rep2, respectively. The Venn diagram shows 33 genes commonly downregulated in both replicates. (D) Gene Ontology (GO) enrichment analysis of the 33 commonly downregulated genes identified in (C). Enriched GO biological process terms are shown, including processes associated with nucleic acid metabolism, RNA processing, and DNA replication initiation. The H4K16ac ChIP-seq data were obtained from a publicly available dataset and were not generated from coffee-treated cells.

### Coffee represses MYC-associated genes enriched for H4K16ac

In our previous study, we demonstrated that coffee treatment suppressed the expression of MYC target genes and suggested that this transcriptional repression contributes to the antiproliferative effects of coffee (Cho *et al*., 2026). We therefore investigated whether the 33 H4K16ac-enriched genes consistently downregulated by coffee treatment (Fig. 3C) included genes associated with MYC signaling. Comparison with the Hallmark MYC Targets V1 and V2 gene sets identified two genes, MCM7 and HNRNPA3, that overlapped with MYC Targets V1 (Fig. 4A). In addition, CDCA7/JPO1 has been reported as a MYC target gene (Prescott *et al*, 2001), whereas NME2 has been implicated in MYC transcriptional regulation (Yao *et al*, 2014). We next examined the H4K16ac enrichment profiles of these four MYC-associated genes using the publicly available ChIP-seq dataset. Genome browser visualization revealed H4K16ac signals across the gene bodies of MCM7, HNRNPA3, CDCA7/JPO1, and NME2 (Fig. 4B). Quantitative analysis confirmed that the mean H4K16ac signals within these gene bodies were approximately 2.5–2.8-fold higher than the genome-wide mean (Fig. 4C). We then examined the expression levels of these genes using our previously generated RNA-seq data. Coffee treatment reduced the expression of all four genes in both RNA-seq replicates (Fig. 4D). Notably, NME2 expression was not detected following coffee treatment in rep2, indicating a marked reduction in its expression. Together, these findings demonstrate that coffee treatment downregulates several MYC-associated genes characterized by H4K16ac enrichment. Considering that coffee treatment also induces H4K16 hypoacetylation and alterations in chromatin structure, these results suggest that changes in chromatin organization may contribute to the repression of MYC-associated genes and, potentially, to the inhibition of MYC-dependent cell proliferation previously reported following coffee treatment (Cho *et al*., 2026).

**Figure 4.**
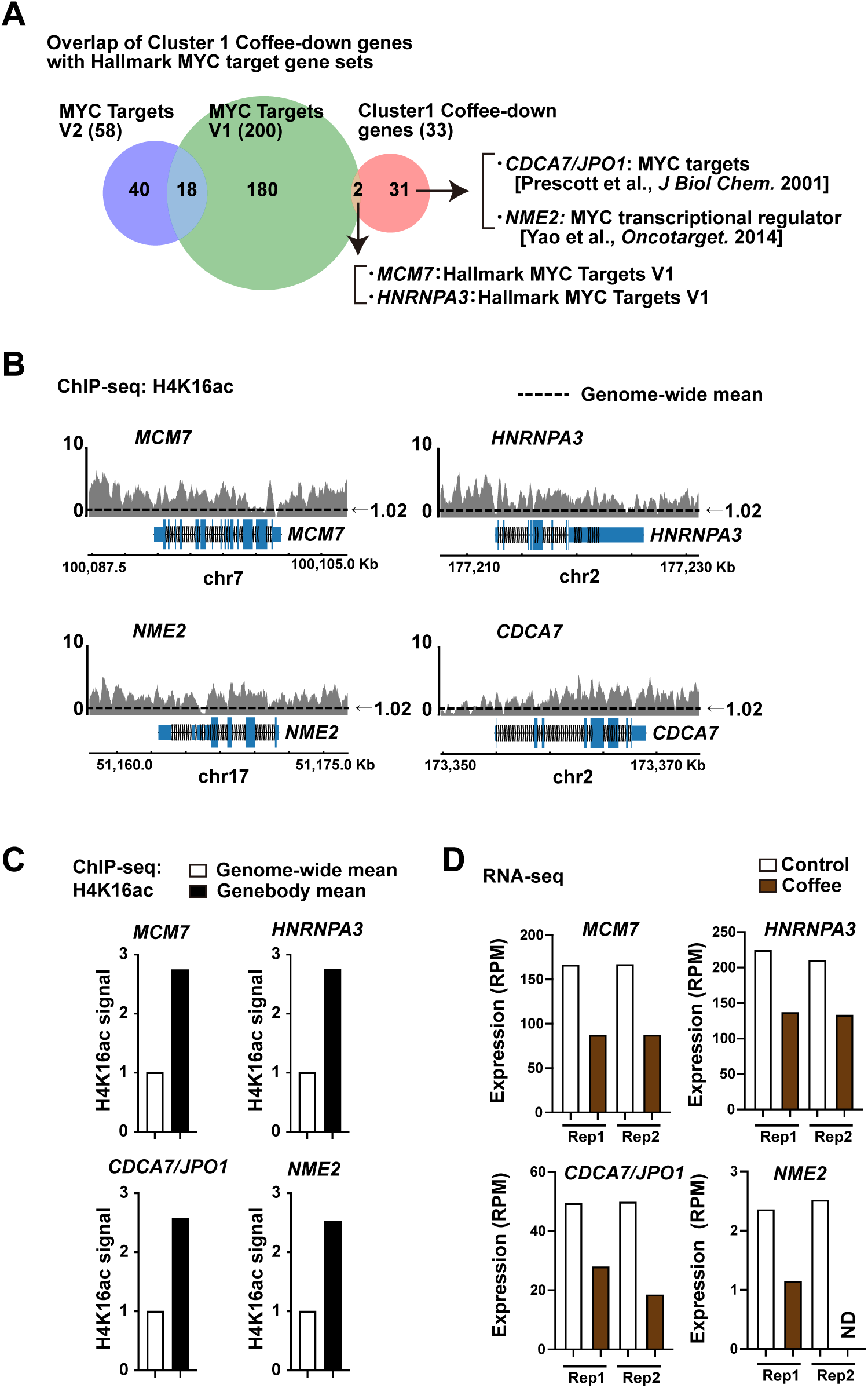
Coffee represses MYC-associated genes enriched for H4K16ac. (A) Identification of MYC-associated genes among the 33 H4K16ac-enriched genes consistently downregulated by coffee treatment. Comparison with the Hallmark MYC Targets V1 and V2 gene sets identified MCM7 and HNRNPA3 as members of MYC Targets V1. CDCA7/JPO1 has previously been reported as a MYC target gene (Prescott et al., 2001), whereas NME2 has been implicated in MYC transcriptional regulation (Yao et al., 2014). (B) Genome browser views showing H4K16ac enrichment at the MCM7, HNRNPA3, CDCA7, and NME2 loci in K562 cells. Signal tracks were obtained from publicly available H4K16ac ChIP-seq data (GSE158736; GSM4809274). (C) Quantification of mean H4K16ac signals across the gene bodies of MCM7, HNRNPA3, CDCA7, and NME2. Gene-body H4K16ac signals were approximately 2.5–2.8-fold higher than the genome-wide mean. (D) RNA-seq expression levels of MCM7, HNRNPA3, CDCA7, and NME2 in control and coffee-treated K562 cells. Expression levels are presented as reads per million (RPM) for two independent biological replicates (rep1 and rep2). Coffee treatment reduced the expression of all four genes in both replicates. NME2 expression was not detected following coffee treatment in rep2. RNA-seq data were obtained from our previous study (Cho et al., 2026), whereas H4K16ac ChIP-seq data were obtained from an independent publicly available dataset.

## Discussion

In the present study, we demonstrated that coffee treatment induces changes in nuclear chromatin organization accompanied by histone H4 lysine 16 hypoacetylation. Western blotting and immunofluorescence analyses confirmed that coffee treatment reduced H4K16ac levels in a cell type-dependent manner (Fig. 1B–E). Notably, HCT116 cells exhibited both reduced H4K16ac levels and increased heterogeneity of nuclear DNA staining intensity following coffee treatment (Figs. 1C–E and 2C). Furthermore, integrative analysis of publicly available H4K16ac ChIP-seq data and previously generated RNA-seq data revealed that a subset of H4K16ac-enriched genes was consistently downregulated by coffee treatment (Fig. 3), including several MYC-associated genes (Fig. 4). Together with our previous findings that coffee suppresses MYC target gene expression and inhibits cell proliferation (Cho *et al*., 2026), these results suggest a potential relationship between coffee-induced epigenetic alterations, chromatin organization, and transcriptional regulation.

Histone H4K16ac plays a particularly important role in the regulation of higher-order chromatin structure. Biochemical studies have demonstrated that H4K16 acetylation disrupts interactions between neighboring nucleosomes and inhibits the formation of compact chromatin fibers(Robinson *et al*., 2008; Shogren-Knaak *et al*., 2006). Furthermore, we previously reported that nuclear c-Abl-mediated signaling induces chromatin structural changes accompanied by H4K16 hypoacetylation(Aoyama *et al*., 2011). In the present study, quantitative analysis of DAPI-stained nuclei revealed a significant increase in the coefficient of variation (CV) of nuclear fluorescence intensity following coffee treatment (Fig. 2C), whereas the mean fluorescence intensity remained unchanged (Fig. 2B). Importantly, this increase in CV was observed in both independent biological replicates, supporting the reproducibility of the observed change in nuclear DNA staining patterns. These findings indicate increased spatial heterogeneity of nuclear DNA staining, consistent with alterations in chromatin organization. Given the established role of H4K16ac in regulating nucleosome interactions, the simultaneous reduction in H4K16ac levels (Fig. 1C–E) raises the possibility that H4K16 hypoacetylation contributes to these structural changes. However, the DAPI-based analysis provides an indirect assessment of chromatin organization, and whether H4K16ac reduction directly causes the observed changes remains to be determined.

An interesting finding of the present study is that coffee-induced H4K16 hypoacetylation varied among cell types. Significant reductions in H4K16ac were observed in K562, HCT116, and A431 cells, whereas changes in MCF7, A549, and TIG-1 cells were not statistically significant (Fig. 1B, C). Notably, our previous study demonstrated that H3K27ac levels were not reduced by coffee treatment in HCT116 cells(Cho *et al*., 2026), despite the significant decrease in H4K16ac observed here. This finding suggests that coffee does not uniformly affect all histone acetylation marks and that its effects depend on both the histone residue and cellular context. H4K16 acetylation is regulated by specific histone acetyltransferases and deacetylases. In particular, MOF/KAT8 is a major acetyltransferase responsible for H4K16 acetylation, whereas sirtuin family deacetylases, including SIRT1, have been implicated in its removal(Smith *et al*, 2005; Taipale *et al*, 2005; Vaquero *et al*, 2004). Thus, differences in the expression, activity, or subcellular localization of histone acetylation regulators may contribute to the cell type-dependent effects of coffee. Furthermore, a mixture of selected coffee constituents examined in our previous study failed to reproduce the reduction in H4K16ac (Fig. 1F), indicating that the specific components responsible for this effect remain unidentified. Further studies will be required to determine whether coffee-induced H4K16 hypoacetylation involves changes in the activity of specific histone acetyltransferases or deacetylases.

Our integrative genomic analysis provided additional insight into the potential relationship between H4K16ac and transcriptional regulation following coffee treatment. H4K16ac has been implicated not only in chromatin organization but also in transcriptional regulation, and genome-wide studies have demonstrated associations between H4K16 acetylation and transcriptionally active chromatin(Bell *et al*, 2007; Feller *et al*, 2012). Analysis of publicly available H4K16ac ChIP-seq data identified four clusters of genes with distinct H4K16ac enrichment patterns (Fig. 3A). Interestingly, genes in Cluster 1 exhibited significantly lower expression fold changes following coffee treatment than those in Clusters 3 and 4 in both RNA-seq replicates (Fig. 3B). Among Cluster 1 genes, 33 were consistently downregulated by more than 1.5-fold in both replicates (Fig. 3C). GO enrichment analysis revealed that these genes were associated with processes involving nucleic acid metabolism, RNA processing, and DNA replication initiation (Fig. 3D). These findings suggest that genes characterized by particular H4K16ac enrichment patterns may be associated with transcriptional repression following coffee treatment. Given the established role of H4K16ac in chromatin organization and transcriptional regulation, it is possible that coffee-induced epigenetic changes contribute to these transcriptional responses. However, because the H4K16ac ChIP-seq dataset was obtained independently of coffee treatment, the present analysis does not establish whether H4K16ac occupancy at these genes is directly altered by coffee.

The potential involvement of MYC-associated genes is particularly relevant to the biological effects of coffee. MYC is a transcription factor that regulates diverse cellular processes, including cell growth, proliferation, metabolism, and ribosome biogenesis(Dang, 2012). Previous studies have also demonstrated functional interactions between MYC and chromatin-modifying factors, including histone acetyltransferase complexes, suggesting that chromatin acetylation contributes to MYC-dependent transcriptional regulation(McMahon *et al*, 2000). In our previous study, we demonstrated that coffee treatment suppresses MYC target gene expression and inhibits cell proliferation(Cho *et al*., 2026). In the present study, we identified four MYC-associated genes, MCM7, HNRNPA3, CDCA7/JPO1, and NME2, among the H4K16ac-enriched genes consistently downregulated by coffee treatment (Fig. 4A). These genes exhibited substantial H4K16ac enrichment across their gene bodies (Fig. 4B, C) and reduced expression following coffee treatment (Fig. 4D). MCM7 is involved in DNA replication, HNRNPA3 participates in RNA processing, CDCA7/JPO1 is a MYC-responsive gene(Prescott *et al*., 2001), and NME2 has been implicated in MYC transcriptional regulation (Yao *et al*., 2014). Thus, repression of these genes is consistent with the suppression of MYC-associated cellular functions observed in our previous study. These findings raise the possibility that coffee-induced alterations in chromatin organization and histone acetylation are associated with the repression of MYC-related transcriptional programs. However, whether H4K16 hypoacetylation directly contributes to the suppression of these genes or to the antiproliferative effects of coffee remains to be established.

Several limitations of the present study should be considered. First, although H4K16 hypoacetylation and changes in nuclear DNA staining patterns were both observed following coffee treatment in HCT116 cells (Figs. 1C–E and 2), their causal relationship has not been established. Second, chromatin structural changes were evaluated using the spatial distribution of nuclear DNA fluorescence intensity, which provides an indirect measure of chromatin organization rather than a direct assessment of higher-order chromatin architecture. Third, the chromatin imaging experiments were performed in HCT116 cells, whereas the integrative ChIP-seq and RNA-seq analyses were conducted using K562 datasets (Figs. 3 and 4). Therefore, the relationship between chromatin structural changes and transcriptional repression has not been directly demonstrated within the same cellular system. In addition, although cell-level differences in nuclear H4K16ac intensity and DAPI fluorescence heterogeneity were observed in both independent experiments, additional biological replicates would strengthen the assessment of these effects across independently treated cultures. Finally, the specific coffee constituents responsible for H4K16 hypoacetylation remain unidentified (Fig. 1F). Future studies involving perturbation of H4K16 acetylation regulators, locus-specific analysis of H4K16ac following coffee treatment, and direct assessment of chromatin accessibility or higher-order organization will be important for clarifying the underlying mechanisms.

In conclusion, the present study demonstrates that coffee treatment reduces H4K16ac levels and alters the spatial distribution of nuclear DNA staining intensity in a cell type-dependent manner (Figs. 1 and 2). Furthermore, integrative genomic analysis revealed that a subset of H4K16ac-enriched genes, including MYC-associated genes, exhibits reproducible transcriptional downregulation following coffee treatment (Figs. 3 and 4). Together with our previous findings on the suppression of MYC target genes and cell proliferation (Cho *et al*., 2026), these results suggest a potential association between coffee-induced H4K16 hypoacetylation, chromatin organization, and transcriptional regulation. Further investigation will be required to establish whether these changes are mechanistically linked.

## Acknowledgments

We thank Dr. Noritaka Yamaguchi (Meiji Pharmaceutical University) for kindly providing the MCF-7, A549, A431, and HCT116 cell lines used in this study. The authors thank Dr. Megumi Tago and Dr. Yosuke Nakazawa (Keio University) for kindly providing the K562 cell line and for general support. The authors thank Junko Yamada for maintaining the shared equipment platform and for her logistical support. This work was supported in part by JSPS KAKENHI (Grant Number 23K07851); the Kobayashi Foundation; the Friends of Leukemia Research Fund (Takaku Fumimaro Award); the Japanese Society of Hematology; the Keio University Academic Development Fund; the Keio University Fukuzawa Fund; the Chemo-Sero-Therapeutic Research Institute (Kaketsuken); the Suzuken Memorial Foundation; the Mochida Memorial Foundation; and the Takeda Science Foundation. ChatGPT (OpenAI) was used under author supervision for minor revisions, consistency checking, and idea exploration. All content was verified by the authors.

## Author Contributions

Y.T. and Y.C. performed the major experiments, contributed to the experimental methodology, and collected and analyzed the data. Y.S., S.M., and R.U. performed the experiments. K.A. conceived and designed the study, performed experiments, analyzed the data, prepared the figures, supervised the research, and wrote the manuscript. All authors reviewed and approved the final manuscript.

## References

Aoyama K, Fukumoto Y, Ishibashi K, Kubota S, Morinaga T, Horiike Y, Yuki R, Takahashi A, Nakayama Y, Yamaguchi N (2011) Nuclear c-Abl-mediated tyrosine phosphorylation induces chromatin structural changes through histone modifications that include H4K16 hypoacetylation. Experimental cell research 317: 2874–2903

Aoyama K, Oshima M, Koide S, Suzuki E, Mochizuki-Kashio M, Kato Y, Tara S, Shinoda D, Hiura N, Nakajima-Takagi Y *t* (2018) Ezh1 Targets Bivalent Genes to Maintain Self-Renewing Stem Cells in Ezh2-Insufficient Myelodysplastic Syndrome. iScience 9: 161–174

Aoyama K, Shinoda D, Suzuki E, Nakajima-Takagi Y, Oshima M, Koide S, Rizq O, Si S, Tara S, Sashida G et al (2021) PRC2 insufficiency causes p53-dependent dyserythropoiesis in myelodysplastic syndrome. Leukemia 35: 1156–1165

Aoyama K, Yamaguchi N, Yuki R, Morii M, Kubota S, Hirata K, Abe K, Honda T, Kuga T, Hashimoto Y et al (2015) c-Abl induces stabilization of histone deacetylase 1 (HDAC1) in a kinase activity-dependent manner. Cell biology international 39: 446–456

Aoyama K, Yuki R, Horiike Y, Kubota S, Yamaguchi N, Morii M, Ishibashi K, Nakayama Y, Kuga T, Hashimoto Y et al (2013) Formation of long and winding nuclear F-actin bundles by nuclear c-Abl tyrosine kinase. Experimental cell research 319: 3251–3268

Bannister AJ, Kouzarides T (2011) Regulation of chromatin by histone modifications. Cell Res 21: 381–395

Bell O, Wirbelauer C, Hild M, Scharf AN, Schwaiger M, MacAlpine DM, Zilbermann F, van Leeuwen F, Bell SP, Imhof A et al (2007) Localized H3K36 methylation states define histone H4K16 acetylation during transcriptional elongation in Drosophila. Embo j 26: 4974–4984

Cho Y, Miyota-Yui M, Nakajima-Takagi Y, Yokomizo T, Oshima M, Iwama A, Aoyama K (2026) Coffee remodels the global histone acetylation landscape with reduced H3K27ac at the *MYC* promoter. bioRxiv: 2026.2008.2001.742200

Dang CV (2012) MYC on the path to cancer. Cell 149: 22–35

Feller C, Prestel M, Hartmann H, Straub T, Söding J, Becker PB (2012) The MOF-containing NSL complex associates globally with housekeeping genes, but activates only a defined subset. Nucleic Acids Res 40: 1509–1522

Fraga MF, Ballestar E, Villar-Garea A, Boix-Chornet M, Espada J, Schotta G, Bonaldi T, Haydon C, Ropero S, Petrie K et al (2005) Loss of acetylation at Lys16 and trimethylation at Lys20 of histone H4 is a common hallmark of human cancer. Nat Genet 37: 391–400

Higashi K, Cho Y, Aoyama K (2026) Multifaceted MRVI1 Serves as a Tumor Suppressor in HCT116 Colorectal Cancer Cells. Biol Pharm Bull 49: 589–593

Higashi K, Tanaka Y, Kosako H, Aoyama K (2025) Identification of MRVI1-Interacting Proteins by Biotin-Based Proximity Labeling Reveals NPM-ALK-Dependent Interaction Dynamics. J Biochem

Kouzarides T (2007) Chromatin modifications and their function. Cell 128: 693–705

Kubota S, Fukumoto Y, Aoyama K, Ishibashi K, Yuki R, Morinaga T, Honda T, Yamaguchi N, Kuga T, Tomonaga T et al (2013) Phosphorylation of KRAB-associated protein 1 (KAP1) at Tyr-449, Tyr-458, and Tyr-517 by nuclear tyrosine kinases inhibits the association of KAP1 and Heterochromatin Protein 1 α(HP1α) with heterochromatin. Journal of Biological Chemistry 288: 17871–17883

Kubota S, Morii M, Yuki R, Yamaguchi N, Yamaguchi H, Aoyama K, Kuga T, Tomonaga T, Yamaguchi N (2015) Role for Tyrosine Phosphorylation of A-kinase Anchoring Protein 8 (AKAP8) in Its Dissociation from Chromatin and the Nuclear Matrix. The Journal of biological chemistry 290: 10891–10904

Mashimo M, Bu X, Aoyama K, Kato J, Ishiwata-Endo H, Stevens LA, Kasamatsu A, Wolfe LA, Toro C, Adams D et al (2019) PARP1 inhibition alleviates injury in ARH3-deficient mice and human cells. JCI insight 4

McMahon SB, Wood MA, Cole MD (2000) The essential cofactor TRRAP recruits the histone acetyltransferase hGCN5 to c-Myc. Molecular and cellular biology 20: 556–562

Prescott JE, Osthus RC, Lee LA, Lewis BC, Shim H, Barrett JF, Guo Q, Hawkins AL, Griffin CA, Dang CV (2001) A novel c-Myc-responsive gene, JPO1, participates in neoplastic transformation. The Journal of biological chemistry 276: 48276–48284

Ramírez F, Ryan DP, Grüning B, Bhardwaj V, Kilpert F, Richter AS, Heyne S, Dündar F, Manke T (2016) deepTools2: a next generation web server for deep-sequencing data analysis. Nucleic Acids Res 44: W160–165

Raudvere U, Kolberg L, Kuzmin I, Arak T, Adler P, Peterson H, Vilo J (2019) g:Profiler: a web server for functional enrichment analysis and conversions of gene lists (2019 update). Nucleic Acids Research 47: W191–W198

Robinson PJ, An W, Routh A, Martino F, Chapman L, Roeder RG, Rhodes D (2008) 30 nm chromatin fibre decompaction requires both H4-K16 acetylation and linker histone eviction. J Mol Biol 381: 816–825

Roth SY, Denu JM, Allis CD (2001) Histone acetyltransferases. Annual review of biochemistry 70: 81–120

Shogren-Knaak M, Ishii H, Sun JM, Pazin MJ, Davie JR, Peterson CL (2006) Histone H4-K16 acetylation controls chromatin structure and protein interactions. *Science (New York*, NY) 311: 844–847

Smith ER, Cayrou C, Huang R, Lane WS, Côté J, Lucchesi JC (2005) A human protein complex homologous to the Drosophila MSL complex is responsible for the majority of histone H4 acetylation at lysine 16. Molecular and cellular biology 25: 9175–9188

Taguchi S, Higashi K, Tanaka Y, Kosako H, Aoyama K (2026) Oncogenic NPM-ALK reprograms the TGM1 interactome toward oncogenic signaling and transcriptional states. bioRxiv: 2026.2008.2001.742203

Taipale M, Rea S, Richter K, Vilar A, Lichter P, Imhof A, Akhtar A (2005) hMOF histone acetyltransferase is required for histone H4 lysine 16 acetylation in mammalian cells. Molecular and cellular biology 25: 6798–6810

Takahashi A, Obata Y, Fukumoto Y, Nakayama Y, Kasahara K, Kuga T, Higashiyama Y, Saito T, Yokoyama KK, Yamaguchi N (2009) Nuclear localization of Src-family tyrosine kinases is required for growth factor-induced euchromatinization. Experimental cell research 315: 1117–1141

Vaquero A, Scher M, Lee D, Erdjument-Bromage H, Tempst P, Reinberg D (2004) Human SirT1 interacts with histone H1 and promotes formation of facultative heterochromatin. Mol Cell 16: 93–105

Yao Y, Li C, Zhou X, Zhang Y, Lu Y, Chen J, Zheng X, Tao D, Liu Y, Ma Y (2014) PIWIL2 induces c-Myc expression by interacting with NME2 and regulates c-Myc-mediated tumor cell proliferation. Oncotarget 5: 8466–8477

Yuki R, Aoyama K, Kubota S, Yamaguchi N, Kubota S, Hasegawa H, Morii M, Huang X, Liu K, Williams R et al (2015) Overexpression of zinc-finger protein 777 (ZNF777) inhibits proliferation at low cell density through down-regulation of FAM129A. J Cell Biochem 116: 954–968

Yuki R, Tatewaki T, Yamaguchi N, Aoyama K, Honda T, Kubota S, Morii M, Manabe I, Kuga T, Tomonaga T et al (2019) Desuppression of TGF-β signaling via nuclear c-Abl-mediated phosphorylation of TIF1γ/TRIM33 at Tyr-524, -610, and -1048. Oncogene 38: 637–655

